# DISPERSE – A trait database to assess the dispersal potential of aquatic macroinvertebrates

**DOI:** 10.1101/2020.02.21.953737

**Authors:** Romain Sarremejane, Núria Cid, Rachel Stubbington, Thibault Datry, Maria Alp, Miguel Cañedo-Argüelles, Adolfo Cordero-Rivera, Zoltán Csabai, Cayetano Gutiérrez-Cánovas, Jani Heino, Maxence Forcellini, Andrés Millán, Amael Paillex, Petr Pařil, Marek Polášek, José Manuel Tierno de Figueroa, Philippe Usseglio-Polatera, Carmen Zamora-Muñoz, Núria Bonada

## Abstract

Dispersal is an essential process in population and community dynamics, but is difficult to measure in the field. In freshwater ecosystems, information on biological traits related to organisms’ morphology, life history and behaviour provides useful dispersal proxies, but information remains scattered or unpublished for many taxa. We compiled information on multiple dispersal-related biological traits of European aquatic macroinvertebrates in a unique resource, the DISPERSE database. DISPERSE includes 39 trait categories grouped into nine dispersal-related traits for 480 taxa, including Annelida, Mollusca, Platyhelminthes, and Arthropoda such as Crustacea and Insecta, generally at the genus level. Information within DISPERSE can be used to address fundamental questions in metapopulation ecology, metacommunity ecology, macroecology and evolutionary ecology research. Information on dispersal proxies can be applied to improve predictions of ecological responses to global change, and to inform improvements to biomonitoring and conservation management strategies. The diverse sources used in DISPERSE complement existing trait databases by providing new information on dispersal traits, most of which would not otherwise be accessible to the scientific community.

## Background & Summary

Dispersal is a fundamental ecological process that affects the organization of biological diversity at multiple temporal and spatial scales^1,2^. Dispersal strongly influences metapopulation and metacommunity dynamics through the movement of individuals and species, respectively^3^. A better understanding of dispersal processes can inform biodiversity management practices^4,5^. However, dispersal is difficult to measure directly, particularly for small organisms, including most invertebrates^6^. Typically, dispersal is measured for single species^7,8^ or combinations of few species within one taxonomic group^9–11^, using methods based on mark and recapture, stable isotopes, or population genetics^5,12^. Such methods can directly assess dispersal events but are expensive, time-consuming, and thus impractical for studies conducted at the community level or at large spatial scales. In this context, taxon-specific biological traits represent a cost-effective alternative that may serve as proxies for dispersal^5,6,13,14^. These traits interact with landscape structure to determine patterns of effective dispersal^15,16^.

Aquatic macroinvertebrates inhabiting freshwater ecosystems include taxa with diverse dispersal modes and abilities (Figure 1). For species with complex life cycles, such as some insects, this diversity is enhanced by distinct dispersal strategies among life stages. For example, juveniles of many aquatic insects disperse actively and/or passively in water whereas adults fly over land^17^. In any case, dispersal is affected by multiple traits relating to the morphology^6,12^, life history and behaviour^2^ of different life stages.

**Figure 1.**
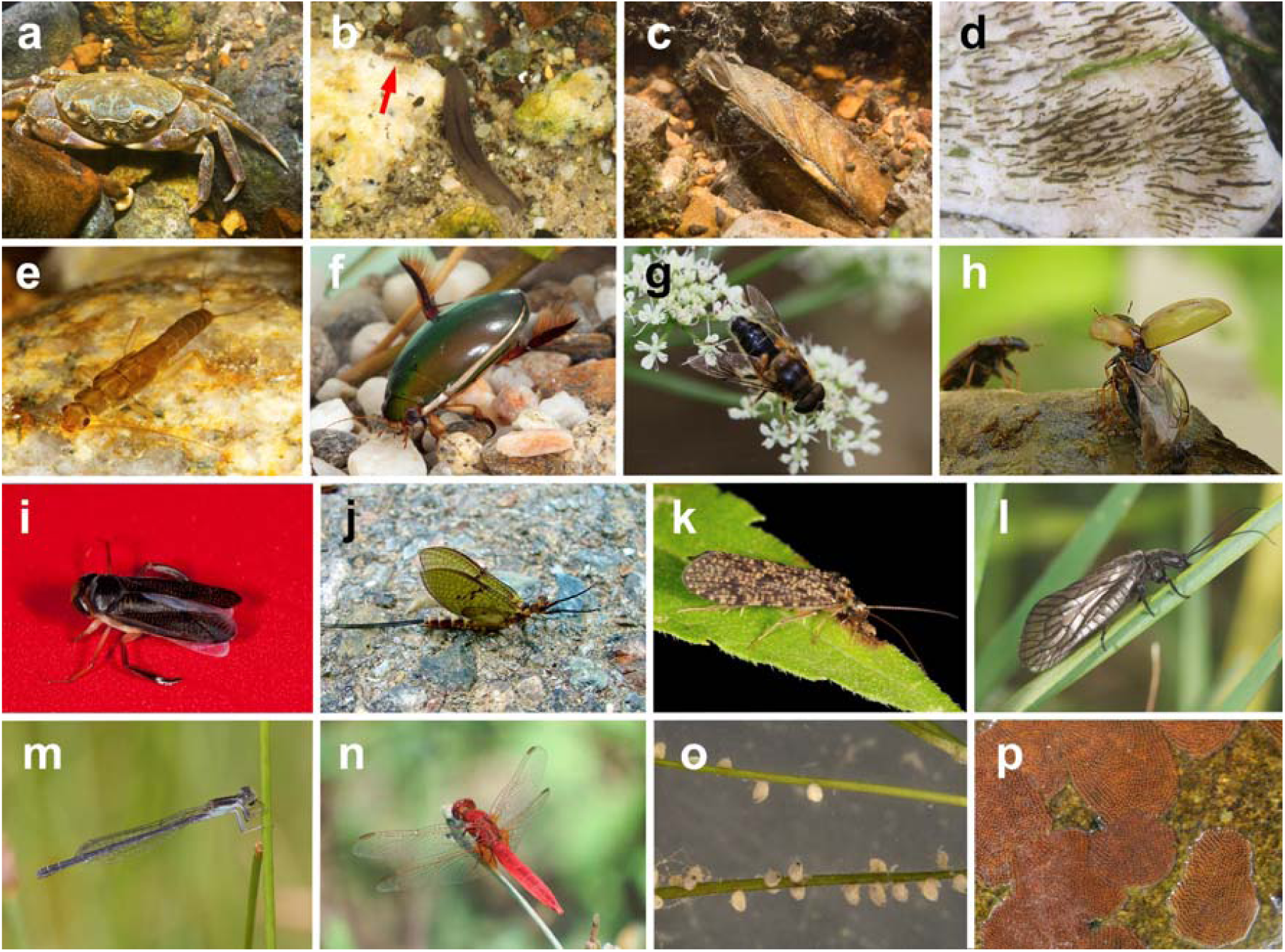
The dispersal-related trait diversity of aquatic macroinvertebrates. Taxa that disperse in water include the crustaceans *Potamon* (a) and *Asellus* (red arrow in b), planarians (b), the bivalve mollusc *Unio* (c), insect larvae such as the Diptera genus *Simulium* (d) and Plecoptera genus *Leuctra* (e), and adult Coleoptera including dytiscid *Cybister* (f). Such aquatic dispersers may move passively in the drift (c, d) and/or actively crawl or swim (a, b, e, f). Most adult insects have wings and can fly overland (f-n). Wings are morphologically diverse and include various types: one wing pair, as in Diptera such as the syrphid genus *Eristalis* (g); one pair of wings with elytra for Coleoptera including the genus *Enochrus* (h) or with hemielytra for Heteroptera such as the genus *Hesperocorixa* (i); two wing pairs including one pair of small hind wings for Ephemoptera including the genus *Ephemera* (j); and two pairs of similar-sized wings for species such as the Trichoptera genus *Polycentropus* (k), the Megaloptera genus *Sialis* (I) or the Odonata genera *Ischnura* (m) and *Crocothemis* (n). Wings range in size from a few mm in some Diptera (g) up to more than 3 cm (l-n), with the Odonata exemplifying the large morphologies. Taxa vary in the number of eggs produced per individual, ranging from tens per reproduction cycle for most Coleoptera and Heteroptera such as the genus *Sigara* (o) to several hundreds in the egg masses of most Ephemeroptera and Trichoptera, such as those of the genus *Hydropsyche* (p). Credits: Adolfo Cordero-Rivera (a-g, i, k-n), Jesús Arribas (h), Pere Bonada (j), José Antonio Carbonell (o) and Maria Alp (p).

We compiled and harmonized information on dispersal-related traits of aquatic macroinvertebrates from across Europe, including both aquatic and aerial (i.e. flying) stages. Although information on some dispersal-related traits such as body size, reproduction, locomotion and dispersal mode is available in online databases for European^18–20^ and North American taxa^21^, other relevant information is scattered across published literature and unpublished data. Using the expertise of 19 experts, we built a comprehensive database containing nine dispersal-related traits divided into 39 trait categories for 480 European taxa. Dispersal-related traits were selected and their trait categories fuzzy-coded^22^ following a structure comparable to existing databases^23^. Our aim was to provide a single resource facilitating the inclusion of dispersal in ecological research, and to create the basis for a global dispersal database.

## Methods

### Dispersal-related trait selection criteria

We defined dispersal as the unidirectional movement of individuals from one location to another^1^, assuming that population-level dispersal rates depend on both the number of dispersers/propagules and dispersers’ ability to move across the landscape^11,24^.

We selected nine dispersal-related morphological, behavioural and life-history traits (Table 1). Selected morphological traits were maximum body size, female wing length and wing pair type, the latter two relating only to flying adult insects. Body size influences organisms’ dispersal^6^, especially for active dispersers^25^, with larger animals more capable of active dispersal over longer distances (e.g. flying adult dragonflies^6^). Wing morphology, and in particular wing length, is related to the dispersal of flying adult insects^6,26^. Female wing length was selected because females connect and sustain populations through oviposition, thus representing adult insects’ flying and recolonization capacity^27^. Females with larger wings are likely to oviposit farther from their source population^6,10,28^. We also described wing morphology as insect wing types, i.e. one or two pairs of wings, and the presence of halters, elytra or hemielytra, or small hind wings^12^ (Figure 1). Selected life-history traits were adult life span, life-cycle duration, annual number of reproductive cycles and lifelong fecundity. Adult life span and life-cycle duration respectively reflect the adult (i.e. reproductive) and total life duration, with longer-lived animals typically having more dispersal opportunities^13^. The annual number of reproductive cycles and lifelong fecundity assess dispersal capacity based on potential propagule production, with multiple reproductive cycles and abundant eggs increasing the expected number of dispersal events^6^. Dispersal behaviour was represented by a taxon’s predominant dispersal mode (passive and/or active, aquatic and/or aerial), and by propensity to drift, which indicates the frequency of flow-mediated passive downstream dispersal.

**Table 1.**
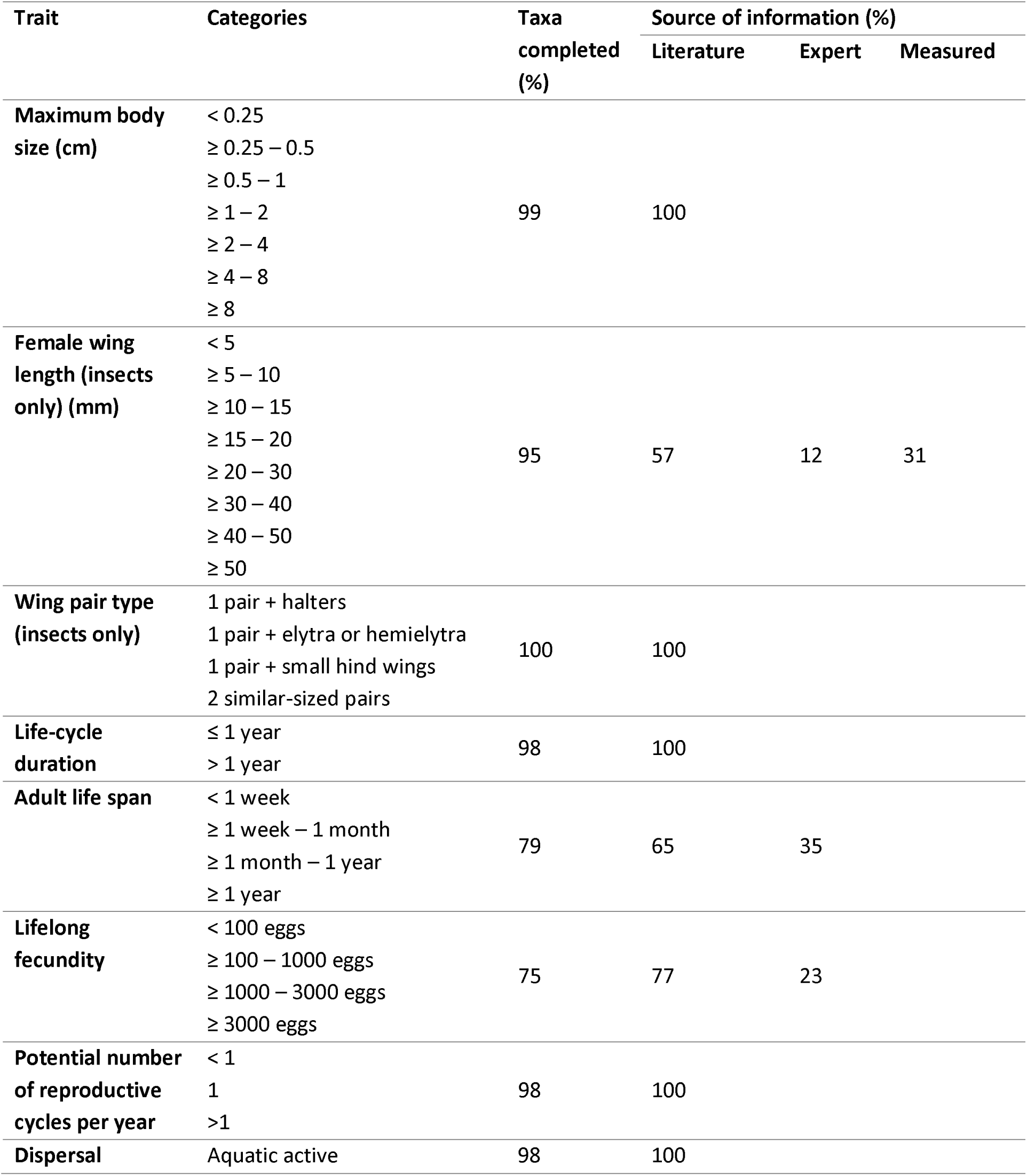
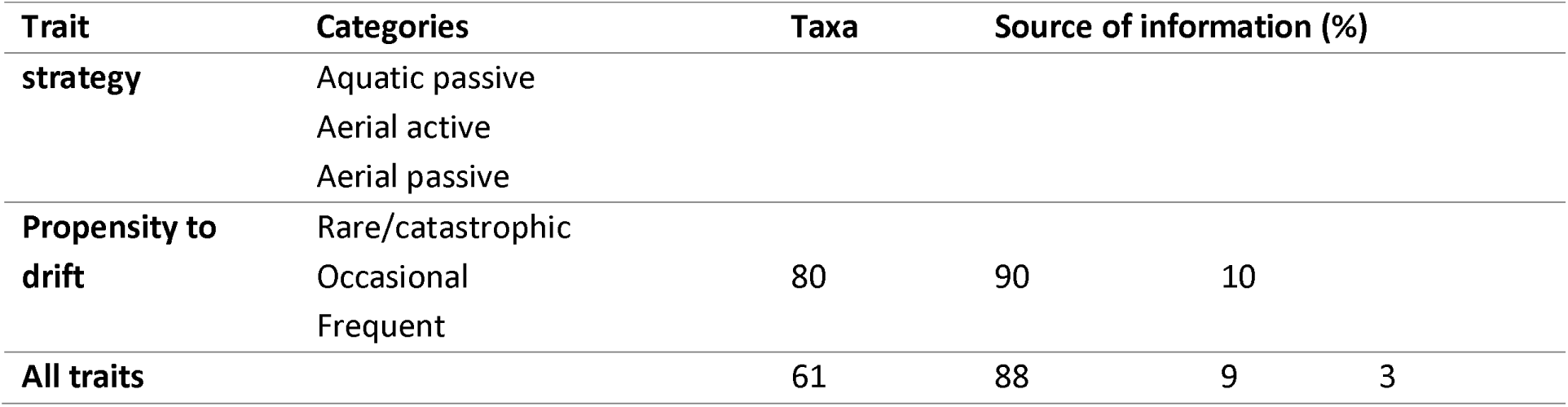
Dispersal-related aquatic macroinvertebrate traits included in the DISPERSE database and the relative contribution of different sources of information (i.e. literature, expert knowledge, direct measurement) used to build the database.

### Data acquisition and compilation

A taxa list was generated based on existing trait databases^20^ and modified to reflect current taxonomic classifications. Trait information was sourced primarily from the literature using Google Scholar searches of keywords including trait names, synonyms and taxon names, and by searching in existing databases^18,21^. Altogether, >300 peer-reviewed articles and book chapters were consulted (Supplementary File 1). When no European studies were available, we considered information from other continents only if experts evaluated traits as comparable across regions. When published information was lacking, traits were coded based on authors’ expert knowledge and direct measurements. Specifically, for 139 species in 69 genera of Coleoptera and Heteroptera, female wing lengths were characterized using measurements of 538 individuals in experts’ reference collections, sampled in Finland, Hungary and Greece. The number of species measured within a genus varied between 1 and 10 in proportion to the number of species within each genus in Europe. For example, for the most specious genera, common and rare species as well as northern latitude and Mediterranean species were included.

### Fuzzy-coding approach and taxonomic resolution

Traits were coded using a fuzzy approach in which a value given to each trait category indicates if the taxon has no (0), weak (1), moderate (2) or strong (3) affinity with the category^22^. This method can incorporate intra-taxon variability when trait profiles differ among e.g. species within a genus, early and late instars, or within one species in different environments^29^. Most traits were coded at genus level, but some Diptera and Annelida were coded at family, sub-families or tribe level because of their complex taxonomy, identification difficulties and the scarcity of reliable information about their traits.

## Data Records

DISPERSE can be downloaded as an Excel spreadsheet from the Intermittent River Biodiversity Analysis and Synthesis webpage^30^ or Figshare at: https://figshare.com/s/4f074fb4bdbbe8cf149e.

The database comprises three sheets: DataKey, Data and Reference list. The “Datakey” sheet summarizes the content of each column in the “Data” sheet. The “Data” sheet includes the fuzzy-coded trait categories and cites the sources used to code each trait. The four first columns list the taxa and their taxonomy (group, family, genus [or lowest taxonomic resolution achieved including e.g. tribes and sub-families] and genus synonyms) to allow users to sort and compile information. Sources are cited in chronological order by the surname of the first author and the year of publication. Expert evaluations are reported as “Unpublished” followed by the name of the expert providing the information. Direct measurements are reported as “Direct measurement from” followed by the expert’s name. The “Reference list” sheet contains the references cited in the “Data” sheet, organized in alphabetical order and then by date (also available in Supplementary File 1).

In total, the database contains nine dispersal-related traits divided into 39 trait categories for 480 taxa. Most (78%) taxa are insects, principally Coleoptera and Trichoptera, as these are, together with Diptera, the most diverse orders in freshwater ecosystems^31^. DISPERSE provides complete trait information for 61% of taxa, with 1-2 traits being incomplete for the 39% remaining taxa (Table 1, Figure 2). The traits with the highest percentage of information across taxa were wing pair type and maximum body size, followed by dispersal strategy, life-cycle duration, potential number of reproductive cycles per year, and female wing length (Table 1). The percentage of completed information was lower for two life-history traits: adult lifespan and lifelong fecundity (Table 1).

**Figure 2.**
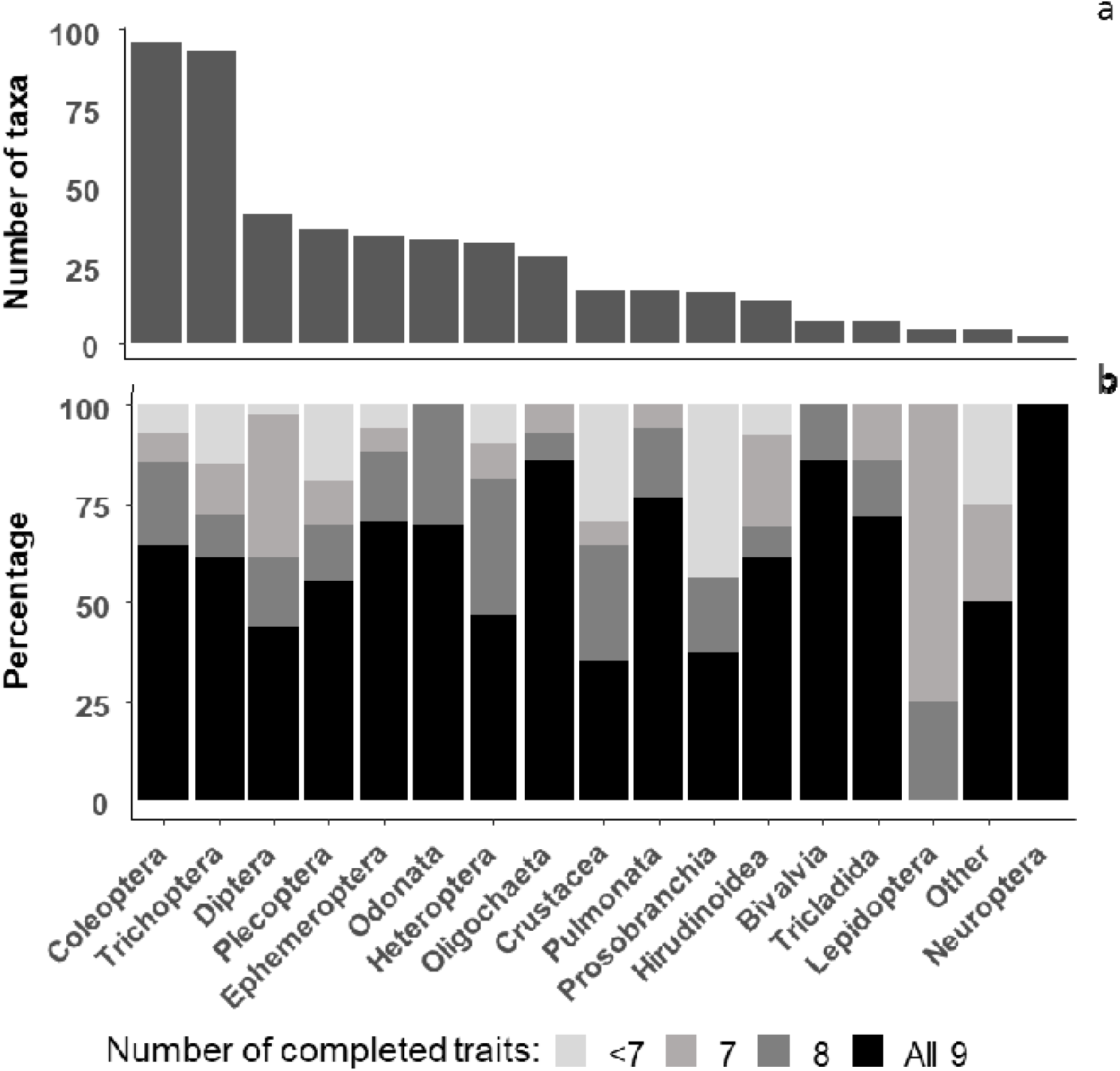
Total number of taxa and percentage of the nine traits completed in each insect order and macroinvertebrate phylum, sub-phylum, class or sub-class. “Other” includes Hydrozoa, Hymenoptera, Megaloptera and Porifera, for which the database includes only one genus.

## Technical Validation

Most of the trait information (88%) originated from published literature (Supplementary File 1) and the rest was coded based on expert knowledge (9%) and direct measurements (3%) (Table 1). The database states information sources for each trait and taxon, allowing users evaluate data quality. Most traits were coded using multiple sources representing multiple species within a genus. When only one study was available, we supplemented this information with expert knowledge, to ensure that trait codes represented potential variability in the taxon.

Using insects as an example, we performed a fuzzy correspondence analysis (FCA)^22^ to visualize variability in trait composition among taxa (Figure 3). Insect orders were clearly distinguished based on their dispersal-related traits, with 32% of the variation explained by the first two FCA axes. Wing pair type and lifelong fecundity had the highest correlation coefficient (0.87 and 0.63, respectively) with axis A1. Female wing length (0.73) and maximum body size (0.55) best correlated with axis A2 (Figures 3 and 4). For example, Coleoptera typically produce few eggs and have intermediate maximum body sizes and wing lengths, Odonata produce an intermediate number of eggs and have long wings, and Ephemeroptera produce many eggs and have short wings (Figures 1 and 4).

**Figure 3.**
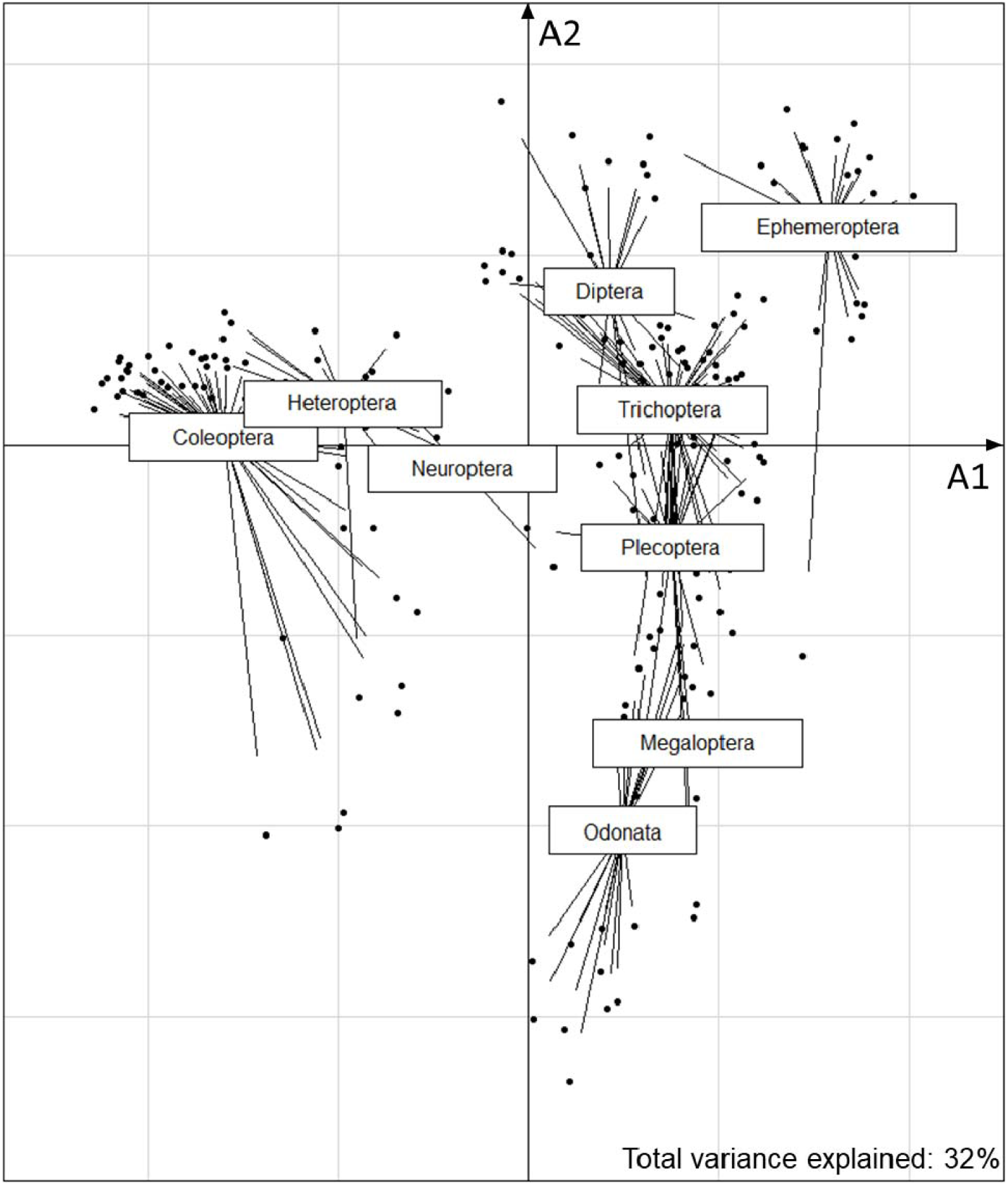
Variability in the dispersal-related trait composition of all insect orders with complete trait profiles along fuzzy correspondence analysis axes A1 and A2. Dots indicate taxa and lines converge to the centroid of each order to depict within-group dispersion.

**Figure 4.**
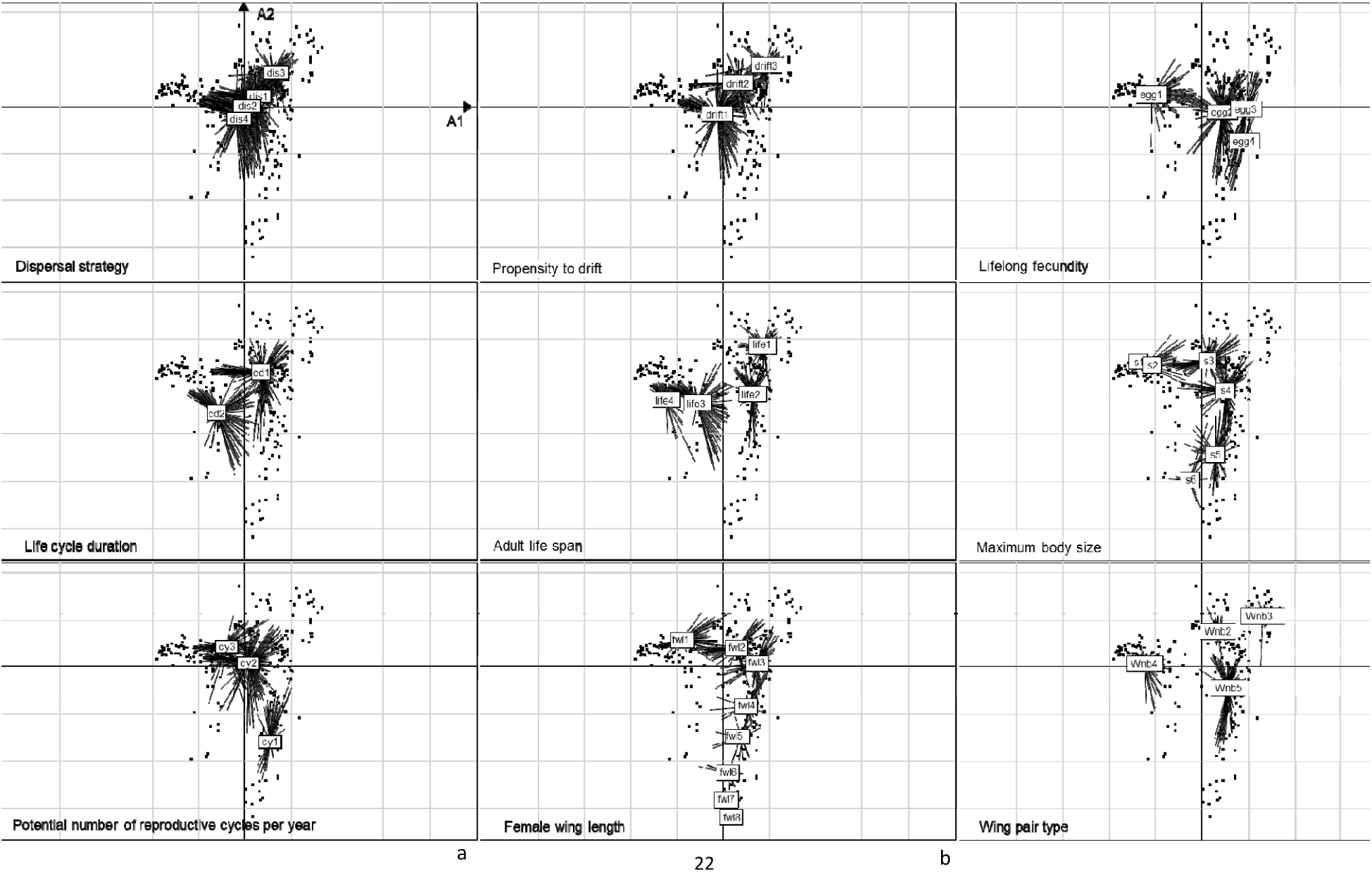
Trait category locations in the fuzzy correspondence analysis ordination space for each trait: (a) Dispersal strategy = **dis1**: aquatic active, **dis2**: aquatic passive, **dis3**: aerial active, **dis4**: aerial passive; (b) Propensity to drift = **drift1**: rare/catastrophic, **drift2**: occasional, **drift3**: frequent; (c) Fecundity = **egg1**: < 100 eggs, **egg2**: ≥ 100 – 1000 **eggs**, egg3: 1000 – 3000 eggs, **egg4**: ≥ 3000 eggs; (d) Life-cycle duration = **cd1**: ≤ 1 year, **cd2**: > 1 year; (e) Adult life span = **life1**: < 1 week, **life2**: ≥ 1 week – 1 month, **life3**: ≥ 1 month – 1 year, l**ife4**: > 1 year; (f) Maximum body size (cm) = **s1**: < 0.25, **s2**: ≥ 0.25 – 0.5, **s3**: ≥ 0.5 – 1, s**4**: ≥ 1 – 2; **s5**: ≥ 2 – 4, **s6**: ≥ 4 – 8; (g) Potential number of reproductive cycles per year = **cy1**: < 1, **cy2**: 1, **cy3**: > 1; (h) Female wing length (cm) = **fwl1**: < 5, **fwl2**: ≥ 5 – 10, **fwl3**: ≥ 10 – 15, **fwl4**: ≥ 15 – 20, **fwl5**: ≥ 20 – 30, **fwl6**: ≥ 30 – 40, **fwl7**: ≥ 40 – 50, **fwl8**: ≥ 50; (i) Wing pair type = Wbn2: 1 pair + halters, Wbn3: 1 pair + elytra or hemielytra, **Wbn4:** 1 pair + small hind wings, **Wbn5**: 2 similar-sized pairs.

The database currently represents a Europe-wide resource, which can be updated and expanded as new information becomes available, to include more taxa and traits from across and beyond Europe. For example, additional information could be collected on other measures of wing morphology^10,14^ and functionality or descriptors of exogenous dispersal vectors such as wind and animals^32^ DISPERSE lays the foundations for a global dispersal trait database, the lack of which is recognized as limiting research progress across multiple disciplines^33^.

## Usage Notes

DISPERSE is the first publicly available database describing the dispersal traits of aquatic macroinvertebrates including information on both aquatic and aerial (i.e. flying) life stages. It provides good coverage of macroinvertebrates at the genus level, which is generally considered as adequate to capture biodiversity dynamics^34–37^. It will promote incorporation of dispersal proxies into fundamental and applied population and community ecology in aquatic ecosystems^5^. In particular, metacommunity ecology may benefit from the use of dispersal traits^15,38^, which enable classification of taxa according to their dispersal potential in more detail. Such classification, used in combination with, for example, spatial distance measurements^39,40^, may advance our understanding of the effects of regional dispersal processes on community assembly and biodiversity patterns. Improved knowledge of taxon-specific dispersal abilities may also inform the design of more effective management practices. For example, recognizing dispersal abilities in biomonitoring methods could inform enhancements to catchment-scale management strategies that support ecosystems adapting to global change^41,42^. DISPERSE could also inform freshwater conservation strategies by establishing different priorities depending on organisms’ dispersal capacities in relation to spatial connectivity^43^.

DISPERSE could also improve species distribution models (SDMs), in which dispersal has rarely been considered due to insufficient data ^13^, limiting the accuracy of model predictions^44^. Recent trait-based approaches have begun to integrate dispersal into SDMs, and DISPERSE could increase model accuracy^45,46^. Including dispersal in SDMs is especially relevant to assessments of biodiversity loss and species vulnerability to climate change^45,47,48^. DISPERSE could also advance understanding of eco-evolutionary relationships and biogeographical phenomena. In an evolutionary context, groups with lower dispersal abilities should be genetically and taxonomically richer due to long-term isolation^49,50^. From a biogeographical perspective, regions affected by glaciations should have species with greater dispersal abilities, enabling postglacial recolonization^51^.

By capturing different dispersal-related biological traits, DISPERSE provides information on organisms’ potential ability to move between localities as well as on recruitment and reproduction^15^. Traits also facilitate comparison of taxa with different dispersal strategies, which could inform studies conducted at large spatial scales, independent of taxonomy^52^.

Users should note that dispersal-related traits included in DISPERSE represent an indirect measure of dispersal, not actual dispersal. Therefore, the database is not intended to substitute population-level studies related to dispersal, but to act as a repository that collates and summarizes information from such studies. As freshwater biodiversity declines at unprecedented rates^53,54^, collecting, harmonizing and sharing dispersal-related data on freshwater organisms will underpin evidence-informed initiatives that seek to support the resilience of ecosystems adapting to global change.

## Acknowledgements

The study was supported by the COST Action CA15113 Science and Management of Intermittent Rivers and Ephemeral Streams (http://www.smires.eu/), funded by COST (European Cooperation in Science and Technology). NC was supported by the French research program Make Our Planet Great Again. MC-A was supported by the MECODISPER project (CTM2017-89295-P) funded by the Spanish Ministerio de Economía, Industria y Competitividad (MINECO) - Agencia Estatal de Investigación (AEI) and cofunded by the European Regional Development Fund (ERDF). AC-R acknowledges funding by MINECO-AEI-ERDF (CGL2014-53140-P). CG-C was supported by the EDRF (COMPETE2020 and PT2020) and the Portuguese Foundation for Science and Technology, through the CBMA strategic program UID/BIA/04050/2019 (POCI-01-0145-FEDER-007569) and the STREAMECO project (Biodiversity and ecosystem functioning under climate change: from the gene to the stream, PTDC/CTA-AMB/31245/2017). PP and MP were supported by the Czech Science Foundation (P505-20-17305S). ZC was supported by the projects EFOP-3.6.1.-16-2016-00004, 20765-3/2018/FEKUTSTRAT and TUDFO/47138/2019-ITM.

## Author contributions

NB, NC and RSa developed the idea and data collection framework. RSa compiled most of the dispersal-related trait data and structured the database. All authors contributed to the addition and checking of information included in the database, and ZC provided direct measurements of several taxa. AC-R designed Figure 1. MA, MC-A, CG-C and MF helped to finalize the database reference list. NC wrote the first draft of the manuscript, and all authors contributed to finalizing it. RSt proofread the manuscript.

## SUPPORTING INFORMATION

**Supplementary File 1.** List of references used to build DISPERSE.

## References

1. Bohonak, A. J. & Jenkins, D. G. Ecological and evolutionary significance of dispersal by freshwater invertebrates. Ecol. Lett. 6, 783–796 (2003).

2. Clobert, J., Baguette, M., Benton, T. G. & Bullock, J. M. Dispersal ecology and evolution. (Oxford Univ. Press, 2012).

3. Heino, J. et al. Metacommunity organisation, spatial extent and dispersal in aquatic systems: Patterns, processes and prospects. Freshw. Biol. 60, 845–869 (2015).

4. Barton, P. S. et al. Guidelines for using movement science to inform biodiversity policy. Environ. Manage. 56, 791–801 (2015).

5. Heino, J. et al. Integrating dispersal proxies in ecological and environmental research in the freshwater realm. Environ. Rev. 25, 334–349 (2017).

6. Rundle, S. D., Bilton, D. T. & Foggo, A. By wind, wings or water: body size, dispersal and range size in aquatic invertebrates. in Body Size: The Structure and Function of Aquatic Ecosystems (eds. Hildrew, A. G., Raffaelli, D. G. & Edmonds-Brown, R.) 186–209 (Cambridge Univ. Press, 2007).

7. Macneale, K. H., Peckarsky, B. L. & Likens, G. E. Stable isotopes identify dispersal patterns of stonefly populations living along stream corridors. Freshw. Biol. 50, 1117–1130 (2005).

8. Troast, D., Suhling, F., Jinguji, H., Sahlén, G. & Ware, J. A global population genetic study of Pantala flavescens. PLoS One 11, 1–13 (2016).

9. French, S. K. & McCauley, S. J. The movement responses of three libellulid dragonfly species to open and closed landscape cover. Insect Conserv. Divers. (2019). doi: 10.1111/icad.12355

10. Arribas, P. et al. Dispersal ability rather than ecological tolerance drives differences in range size between lentic and lotic water beetles (Coleoptera: Hydrophilidae). J. Biogeogr. 39, 984–994 (2012).

11. Lancaster, J. & Downes, B. J. Dispersal traits may reflect dispersal distances, but dispersers may not connect populations demographically. Oecologia 184, 171–182 (2017).

12. Lancaster, J. & Downes, B. J. Dispersal in the terrestrial environment. in Aquatic Entomology (eds. Lancaster, J. & Downes, B. J.) 137–153 (Oxford Univ. Press, 2013).

13. Stevens, V. M. et al. Dispersal syndromes and the use of life-histories to predict dispersal. Evol. Appl. 6, 630–642 (2013).

14. Outomuro, D. & Johansson, F. Wing morphology and migration status, but not body size, habitat or Rapoport’s rule predict range size in North-American dragonflies (Odonata: Libellulidae). Ecography 42, 309–320 (2019).

15. Tonkin, J. D. et al. The role of dispersal in river network metacommunities: patterns, processes, and pathways. Freshw. Biol. 63, 141–163 (2018).

16. Brown, B. L. & Swan, C. M. Dendritic network structure constrains metacommunity properties in riverine ecosystems. J. Anim. Ecol. 79, 571–580 (2010).

17. Wikelski, M. et al. Simple rules guide dragonfly migration. Biol. Lett. 2, 325–329 (2006).

18. Schmidt-Kloiber, A. & Hering, D. www.freshwaterecology.info – An online tool that unifies, standardises and codifies more than 20,000 European freshwater organisms and their ecological preferences. Ecol. Indic. 53, 271–282 (2015).

19. Serra, S. R. Q., Cobo, F., Graça, M. A. S., Dolédec, S. & Feio, M. J. Synthesising the trait information of European Chironomidae (Insecta: Diptera): towards a new database. Ecol. Indic. 61, 282–292 (2016).

20. Tachet, H., Richoux, P., Bournaud, M. & Usseglio-Polatera, P. Invertébrés d’Eau Douce: Systématique, Biologie, écologie. (CNRS éditions, 2010).

21. Vieira, N. K. M. et al. A Database of Lotic Invertebrate Traits for North America. U.S. Geological Survey Data Series 187 (2006).

22. Chevenet, F., Dolédec, S. & Chessel, D. A fuzzy coding approach for the analysis of long-term ecological data. Freshw. Biol. 31, 295–309 (1994).

23. Schmera, D., Podani, J., Heino, J., Erös, T. & Poff, N. L. R. A proposed unified terminology of species traits in stream ecology. Freshw. Sci. 34, 823–830 (2015).

24. Lancaster, J., Downes, B. J. & Arnold, A. Lasting effects of maternal behaviour on the distribution of a dispersive stream insect. J. Anim. Ecol. 80, 1061–1069 (2011).

25. Jenkins, D. G. et al. Does size matter for dispersal distance? Glob. Ecol. Biogeogr. 16, 415–425 (2007).

26. Harrison, R. G. Dispersal polymorphisms in insects. Annu. Rev. Ecol. Syst. 11, 95–118 (1980).

27. Graham, E. S., Storey, R. & Smith, B. Dispersal distances of aquatic insects: upstream crawling by benthic EPT larvae and flight of adult Trichoptera along valley floors. New Zeal. J. Mar. Freshw. Res. 51, 146–164 (2017).

28. Hoffsten, P. O. Site-occupancy in relation to flight-morphology in caddisflies. Freshw. Biol. 49, 810–817 (2004).

29. Bonada, N. & Dolédec, S. Does the Tachet trait database report voltinism variability of aquatic insects between Mediterranean and Scandinavian regions? Aquat. Sci. 80, 1–11 (2018).

30. Sarremejane, R. et al. DISPERSE: a trait database to assess the dispersal potential of aquatic macroinvertebrates. Intermittent River Biodiversity Analysis and Synthesis (2020). Available at: http://irbas.inrae.fr/latest-news/disperse-database-sarremejane-et-al-2020.

31. Lévêque, C., Balian, E. V & Martens, K. An assessment of animal species diversity in continental waters. Hydrobiologia 542, 39–67 (2005).

32. Green, A. J. & Figuerola, J. Recent advances in the study of long-distance dispersal of aquatic invertebrates via birds. Divers. Distrib. 11, 149–156 (2005).

33. Maasri, A. A global and unified trait database for aquatic macroinvertebrates: the missing piece in a global approach. Front. Environ. Sci. 7, 1–3 (2019).

34. Cañedo-Argüelles, M. et al. Dispersal strength determines meta-community structure in a dendritic riverine network. J. Biogeogr. 42, 778–790 (2015).

35. Datry, T. et al. Metacommunity patterns across three Neotropical catchments with varying environmental harshness. Freshw. Biol. 61, 277–292 (2016).

36. Swan, C. M. & Brown, B. L. Metacommunity theory meets restoration: isolation may mediate how ecological communities respond to stream restoration. Ecol. Appl. 27, 2209–2219 (2017).

37. Sarremejane, R., Mykrä, H., Bonada, N., Aroviita, J. & Muotka, T. Habitat connectivity and dispersal ability drive the assembly mechanisms of macroinvertebrate communities in river networks. Freshw. Biol. 62, 1073–1082 (2017).

38. Jacobson, B. & Peres-Neto, P. R. Quantifying and disentangling dispersal in metacommunities: How close have we come? How far is there to go? Landsc. Ecol. 25, 495–507 (2010).

39. Sarremejane, R. et al. Do metacommunities vary through time? Intermittent rivers as model systems. J. Biogeogr. 44, 2752–2763 (2017).

40. Datry, T., Moya, N., Zubieta, J. & Oberdorff, T. Determinants of local and regional communities in intermittent and perennial headwaters of the Bolivian Amazon. Freshw. Biol. 61, 1335–1349 (2016).

41. Cid, N. et al. A metacommunity approach to improve biological assessments in highly dynamic freshwater ecosystems. Bioscience 70, 427–438 (2020).

42. Datry, T., Bonada, N. & Heino, J. Towards understanding the organisation of metacommunities in highly dynamic ecological systems. Oikos 125, 149–159 (2016).

43. Hermoso, V., Cattarino, L., Kennard, M. J., Watts, M. & Linke, S. Catchment zoning for freshwater conservation: refining plans to enhance action on the ground. J. Appl. Ecol. 52, 940–949 (2015).

44. Thuiller, W. et al. A road map for integrating eco-evolutionary processes into biodiversity models. Ecol. Lett. 16, 94–105 (2013).

45. Willis, S. G. et al. Integrating climate change vulnerability assessments from species distribution models and trait-based approaches. Biol. Conserv. 190, 167–178 (2015).

46. Cooper, J. C. & Soberón, J. Creating individual accessible area hypotheses improves stacked species distribution model performance. Glob. Ecol. Biogeogr. 27, 156–165 (2018).

47. Markovic, D. et al. Europe’s freshwater biodiversity under climate change: distribution shifts and conservation needs. Divers. Distrib. 20, 1097–1107 (2014).

48. Bush, A. & Hoskins, A. J. Does dispersal capacity matter for freshwater biodiversity under climate change? Freshw. Biol. 62, 382–396 (2017).

49. Bohonak, A. J. Dispersal, gene flow, and population structure. Q. Rev. Biol. 74, 21–45 (1999).

50. Dijkstra, K.-D. B., Monaghan, M. T. & Pauls, S. U. Freshwater biodiversity and aquatic insect diversification. Annu. Rev. Entomol. 59, 143–163 (2014).

51. Múrria, C. et al. Local environment rather than past climate determines community composition of mountain stream macroinvertebrates across Europe. Mol. Ecol. 26, 6085–6099 (2017).

52. Statzner, B. & Bêche, L. A. Can biological invertebrate traits resolve effects of multiple stressors on running water ecosystems? Freshw. Biol. 55, 80–119 (2010).

53. Strayer, D. L. & Dudgeon, D. Freshwater biodiversity conservation: recent progress and future challenges. J. North Am. Benthol. Soc. 29, 344–358 (2010).

54. Reid, A. J. et al. Emerging threats and persistent conservation challenges for freshwater biodiversity. Biol. Rev. 94, 849–873 (2019).

